# The DNA Methylome of Inflammatory Bowel Disease (IBD) reflects intrinsic and extrinsic factors in intestinal epithelial cells

**DOI:** 10.1101/565200

**Authors:** Iolanda Agliata, Nora Fernandez-Jimenez, Chloe Goldsmith, Julien C. Marie, Jose Ramon Bilbao, Robert Dante, Hector Hernandez-Vargas

**Author notes:** **Correspondence to:** HH.

## Abstract

Abnormal DNA methylation has been described in human inflammatory conditions of the gastrointestinal tract, such as inflammatory bowel disease (IBD). As other complex diseases, IBD results from the balance between genetic predisposition and environmental exposures. As such, DNA methylation may be placed as an effector of both, genetic susceptibility variants and/or environmental signals such as cytokine exposure. We attempted to discern between these two non-excluding possibilities by performing a meta-analysis of DNA methylation data in intestinal epithelial cells of IBD and control samples. We identified abnormal DNA methylation at different levels: deviation from mean methylation signals at site and region levels, and differential variability. A fraction of such changes are associated with genetic polymorphisms linked to IBD susceptibility. In addition, by comparing with another intestinal inflammatory condition (i.e. celiac disease) we propose that aberrant DNA methylation can also be the result of unspecific processes such as chronic inflammation. Our characterization suggests that IBD methylomes combine intrinsic and extrinsic responses in intestinal epithelial cells, and could point to knowledge-based biomarkers of IBD detection and progression.

**Graphical Abstract:** Conceptual representation of the study. Using a meta-analysis strategy we identified differentially methylated positions or regions (DMP/DMR) in IBD. Our assumption is that gene expression changes (IBD phenotype) take place downstream of DNA methylation. In turn, abnormal DNA methylation can be explained by a direct effect of inflammatory cytokines (“signaling”) and/or the result of a genetic polymorphism (SNP). SNP-DMP associations are called methylation quantitative trait loci (mQTL).

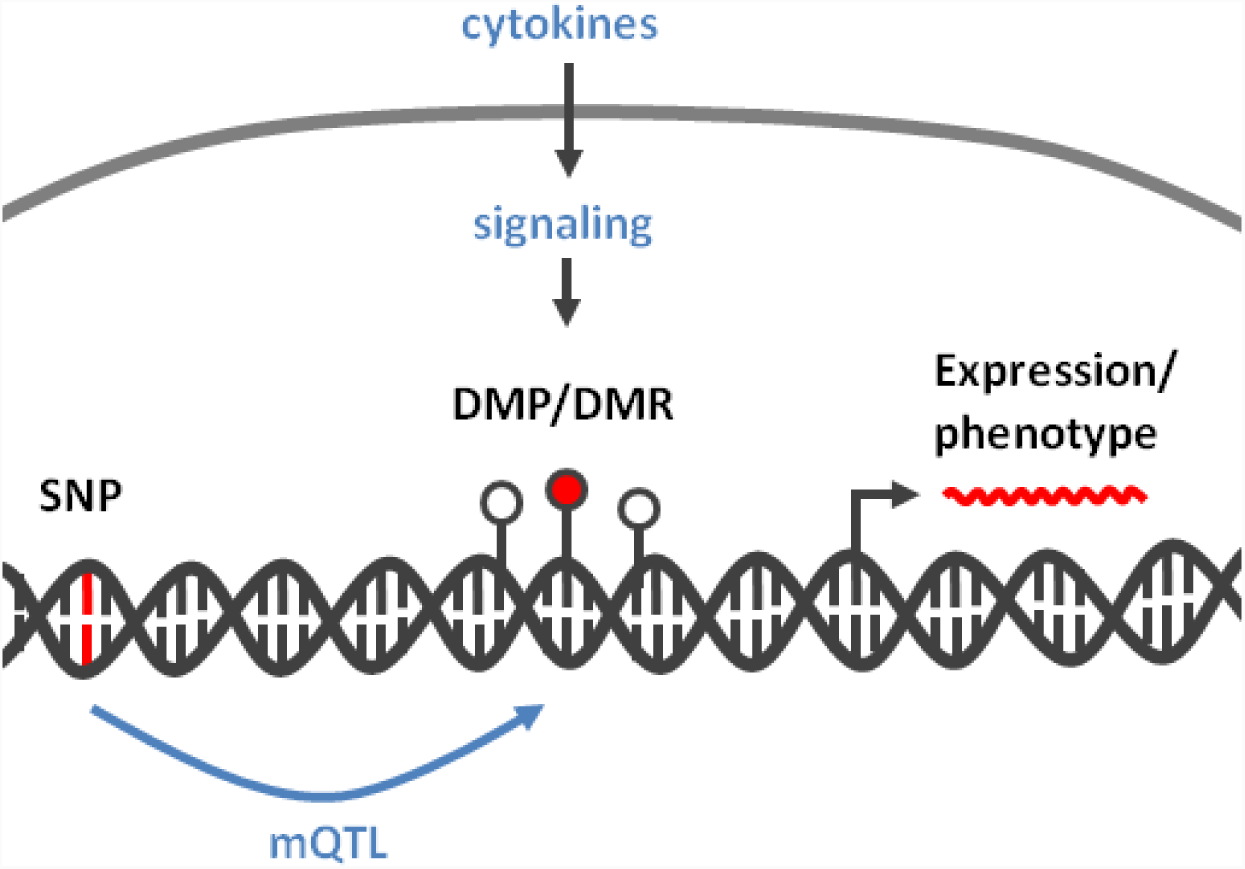

## Background

Inflammatory bowel disease (IBD) comprises Crohn’s disease (CD) and Ulcerative Colitis (UC), two chronic and progressive inflammatory conditions of the gastrointestinal (GI) tract that affect about 2.2 million people in Europe and 1.4 million in United States ^1,2^. The exact etiology is not known, but IBD is characterized by various genetic abnormalities that result in aggressive response from both innate (i.e. macrophages and neutrophils) and acquired (i.e. T and B cells) immunity ^3^. In CD, although inflammation may involve the entire GI tract, the ileum is mainly affected ^4^. In UC, chronic and relapsing inflammation affects the colon and rectum ^5^ and is associated with increased risk of colon cancer development ^6^.

While genetics explains a fraction of inheritance of IBD (13,1% variance in CD and 8,2% in UC) ^7^, environmental factors are able to influence susceptibility through non-genetic mechanisms, such as DNA methylation ^8^. Indeed, several recent studies have provided a detailed characterization of genomic abnormalities in IBD, including DNA methylation ^9–11^. Mechanistically placed between the genome and the transcriptome, DNA methylation may represent an effector of genetic variants and the resulting pathological phenotype ^8^. In addition, DNA methylation is able to perpetuate the response to anti- and pro-inflammatory signals. For example, exposure to cytokines such as interleukin 6 (IL6) and transforming growth factor beta (TGF-β) have been associated with stable DNA methylation changes in epithelial cells ^12–14^. However, it is unclear to what extent the altered DNA methylation of epithelial cells in IBD could be due to persistent cytokine exposure and/or to the direct consequence of genetic susceptibility variants (i.e. SNPs).

Explaining the origin of DNA methylation changes in IBD, may be of interest when exploiting their potential as biomarkers. Currently, the most used biomarkers for IBD are C-Reactive Protein and Calprotectin, although they are not specific for inflammation of intestinal origin, limiting their clinical use ^15^. Instead, DNA methylation is known to be tissue specific ^16,17^, and it may represent a sensor of cytokine exposures ^18–21^ and thus a better biomarker of IBD. Moreover, DNA markers are advantageous in terms of stability, improved isolation and storage, relative to RNA or protein ^22^. With these assumptions, we performed a meta-analysis of intestinal epithelium methylomes in IBD. Our goal was to identify candidate loci that can be potentially useful as biomarkers, using base-resolution methylation data in mucosal biopsies from a large aggregated dataset of CD and UC patients, an approach that may open the way to personalized prevention strategies.

## Results

### Genome-wide changes in DNA methylation are a common feature of IBD

To identify DNA methylation changes in epithelial cells of the intestinal mucosa associated with IBD, we reanalyzed bead-array methylation data from different datasets (**Table 1**). Samples from these datasets included pediatric and adult IBD patients, from both sexes, and involved the two main forms of the condition (i.e. CD and UC).

**Table 1.** Characteristics of the datasets included in the study. PMID: PubMed ID. Idat: raw-level bead-array data availability.

| Accession | Condition | Technique | idat | Samples | Age | Origin | PMID |
| --- | --- | --- | --- | --- | --- | --- | --- |
| MTAB_5463 | UC/CD | HM450/EPIC | yes | 111/104 | 6-15 | Europe | 29031501<sup>9</sup> |
| GSE32146 | UC/CD | HM450 | no | 25 | 14-17 | USA | NA |
| MTAB_3703/3709 | UC/CD | HM450 | yes | 12 | 14 | Europe | 26376367<sup>11</sup> |
| GSE81211 | UC | HM450 | yes | 12 | unknown | South Korea | 27517910<sup>10</sup> |
| GSE105798 | CD | HM450 | yes | 11 | unknown | South Korea | NA |
| GSE42921 | UC/CD | HM450 | no | 23 | 9-16 | USA | NA |

After filtering (see Methods), we tested for the association between IBD and DNA methylation at 393112 CpG sites (81 control and 204 IBD patients) using a linear model. In such a model, we adjusted for sex, age, dataset, and surrogate variables identified during data preprocessing (Fig S1). To account for statistical inflation, we used criteria of effect size (change in mean methylation of at least 10% between controls and IBD) and FDR-adjusted p value < 0.05. Using these criteria, we identified 4280 differentially methylated positions (DMPs), out of which 437 were hypo- and 3843 were hypermethylated in IBD (Fig 1A, Table 2 and Table S1). DMPs were robust to IBD type (Fig 1B), and other clinical and technical features (Fig 1C and S1). An important fraction of these sites were previously identified, in particular in the large dataset published by Howell et al ^9^. However, our dataset combination strategy has led to the identification of new associations. Moreover, the consistency of these findings across independent studies provides additional confidence on their robustness.

**Table 2.**
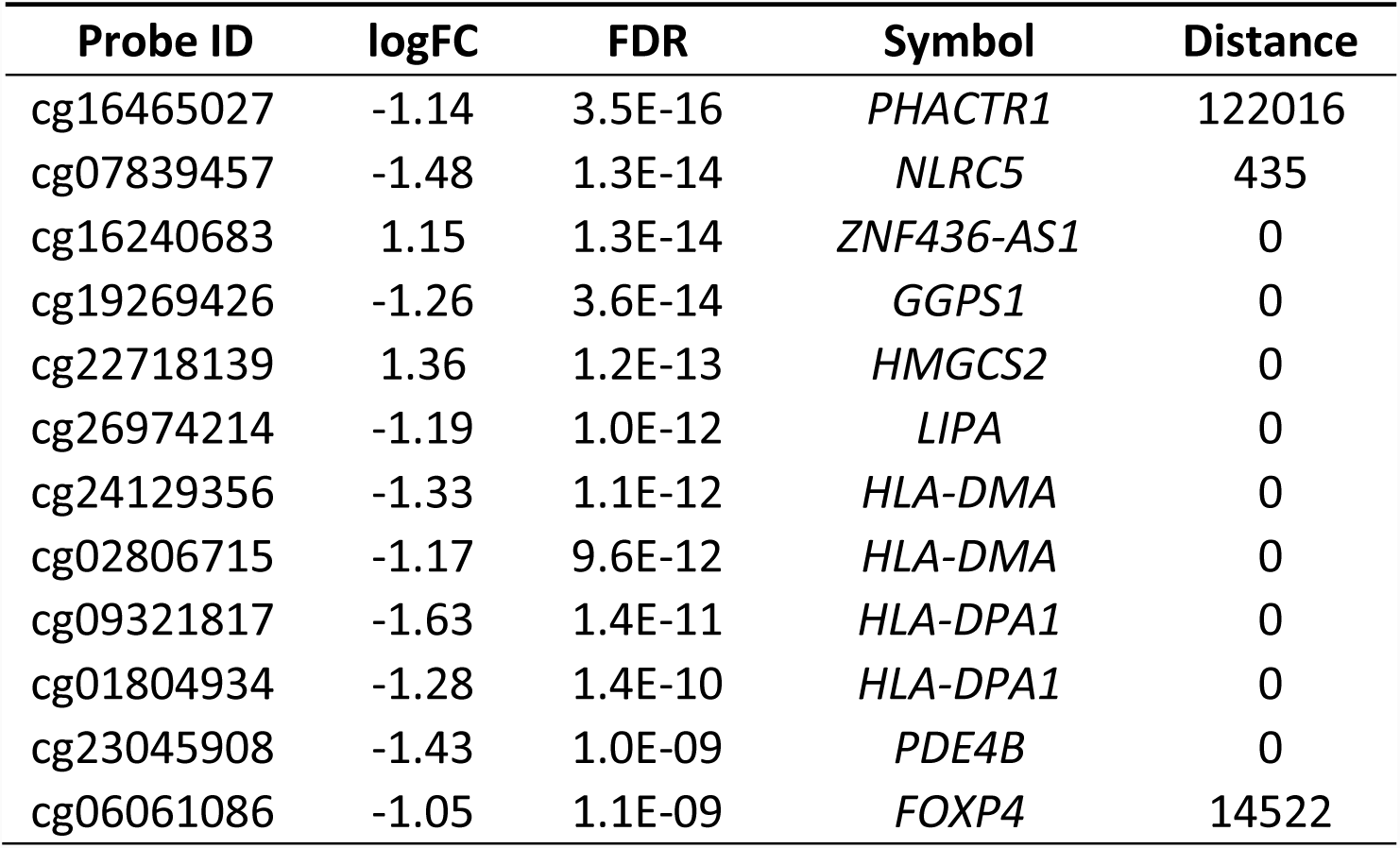
Top DMPs. Differentially methylated positions (DMPs) with a mean difference between groups of at least 20% (delta-beta > 20, FDR < 0.05). Probe ID: Illumina probe reference, logFC: logarithmic fold-change between groups (IBD vs control), FDR: false discovery rate, Symbol: gene symbol, Distance: distance in base pairs to the closest gene. Full list of DMPs can be found in Supplementary Table S1.

| Probe ID | logFC | FDR | Symbol | Distance |
| --- | --- | --- | --- | --- |
| cg16465027 | -1.14 | 3.5E-16 | *PHACTR1* | 122016 |
| cg07839457 | -1.48 | 1.3E-14 | *NLRC5* | 435 |
| cg16240683 | 1.15 | 1.3E-14 | *ZNF436-AS1* | 0 |
| cg19269426 | -1.26 | 3.6E-14 | *GGPS1* | 0 |
| cg22718139 | 1.36 | 1.2E-13 | *HMGCS2* | 0 |
| cg26974214 | -1.19 | 1.0E-12 | *LIPA* | 0 |
| cg24129356 | -1.33 | 1.1E-12 | *HLA-DMA* | 0 |
| cg02806715 | -1.17 | 9.6E-12 | *HLA-DMA* | 0 |
| cg09321817 | -1.63 | 1.4E-11 | *HLA-DPA1* | 0 |
| cg01804934 | -1.28 | 1.4E-10 | *HLA-DPA1* | 0 |
| cg23045908 | -1.43 | 1.0E-09 | *PDE4B* | 0 |
| cg06061086 | -1.05 | 1.1E-09 | *FOXP4* | 14522 |

**Figure 1.**
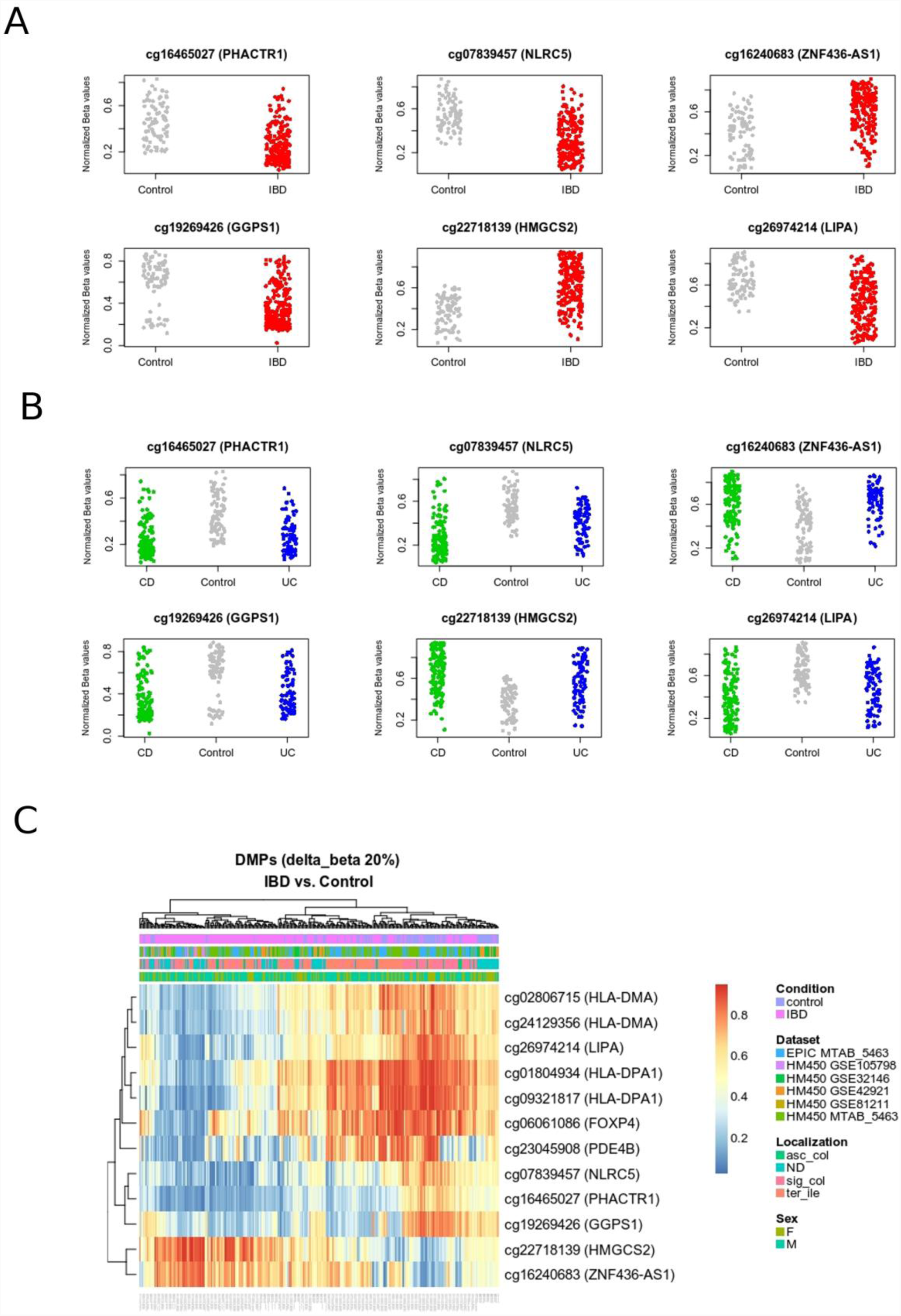
DNA methylation distinguishes IBD from healthy intestinal epithelial cells. **A.** Top differentially methylated positions (DMPs) with a mean difference between IBD (red) vs Control (gray) of at least 20% (delta-beta>20, FDR<0.05). Probe ID and corresponding nearest gene are shown for each significant CpG site. Methylation is represented on the y axis as normalized beta values. **B.** The same CpG sites shown in (A) are represented separately for ulcerative colitis (UC) and Crohn’s disease (CD), shown in blue and green, respectively. **C.** Heatmap showing top differentially methylated positions between IBD *vs* control. The red to blue color gradient represents higher to lower methylation. Main covariates considered in the analysis (i.e. dataset, anatomical location, and sex) are also represented.

An important proportion of DMPs were in the vicinity of each other, suggesting a non-random association with particular genomic loci. To explore this observation, we performed region-level analysis in the same combined dataset. This led to the identification of 1017 differentially methylated regions (DMRs), 172 hypo and 845 hyper methylated in IBD (Tables 3 and S2). As expected, many of these regions corresponded to gene loci also identified using the probe-level strategy (Fig 2B).

**Table 3.** Top DMRs. Differentially methylated regions (DMRs) with at least five CpG sites, a maximum beta change between groups (beta FC) of at least 10%, and a minimum FDR of 0.05, are shown below. # CpGs: number of CpG sites per region, FDR: false discovery rate, Beta FC: methylation beta value fold change. Full list of DMRs can be found in Supplementary Table S2. The table is sorted by beta FC, from hypo- to hypermethylation in IBD.

| Symbol | Coordinates | # CpG | FDR | beta FC |
| --- | --- | --- | --- | --- |
| *HLA-DPA1* | chr6:33040535-33041697 | 8 | 6.5E-87 | -0.216 |
| *VWA1* | chr1:1368846-1370775 | 9 | 9.1E-67 | -0.178 |
| *BST2* | chr19:17516282-17517008 | 5 | 2.2E-83 | -0.155 |
| *HLA-DRA* | chr6:32407289-32408284 | 8 | 4.2E-24 | -0.132 |
| *CIITA* | chr16:10969805-10971250 | 6 | 2.2E-54 | -0.129 |
| *IL12RB1* | chr19:18197544-18198611 | 5 | 1.3E-32 | -0.125 |
| *DAPP1* | chr4:100737138-100738139 | 5 | 1.3E-29 | -0.114 |
| *FCMR* | chr1:207095153-207096833 | 6 | 3.1E-45 | -0.113 |
| *SH3BP2* | chr4:2813458-2814122 | 5 | 2.6E-23 | -0.103 |
| *PRR26* | chr10:695301-696356 | 10 | 2.3E-88 | 0.100 |
| *SPPL2B* | chr19:2278451-2278847 | 5 | 3.4E-62 | 0.100 |
| *HNF1A* | chr12:121415506-121416796 | 8 | 3.4E-105 | 0.100 |
| *FABP1* | chr2:88427027-88428542 | 8 | 5.9E-97 | 0.101 |

| **Symbol** | **Coordinates** | **# CpG** | **FDR** | **beta FC** |
| --- | --- | --- | --- | --- |
| *AATK* | chr17:79107083-79108224 | 7 | 1.6E-37 | 0.101 |
| *HNF4A* | chr20:42983920-42984878 | 12 | 6.2E-140 | 0.101 |
| *LAMA3* | chr18:21452730-21452895 | 6 | 8.0E-74 | 0.102 |
| *MACROD1* | chr11:63852804-63853037 | 5 | 1.8E-28 | 0.102 |
| *QTRT1* | chr19:10822935-10824846 | 7 | 7.7E-75 | 0.102 |
| *LGALS4* | chr19:39303506-39305100 | 5 | 4.0E-38 | 0.102 |
| *RASAL3* | chr19:15568360-15568935 | 5 | 7.6E-62 | 0.102 |
| *LINC00982* | chr1:2979311-2980937 | 8 | 2.3E-88 | 0.103 |
| *JAK3* | chr19:17942221-17943313 | 5 | 3.4E-47 | 0.103 |
| *BTNL8* | chr5:180325254-180326186 | 5 | 1.2E-48 | 0.104 |
| *LINC02372* | chr12:127544951-127545433 | 6 | 1.8E-32 | 0.105 |
| *TMED6* | chr16:69385547-69386847 | 6 | 7.5E-58 | 0.105 |
| *ELMO3* | chr16:67231928-67233983 | 9 | 6.2E-89 | 0.106 |
| *RNU5F-1* | chr1:220132091-220132728 | 6 | 5.4E-53 | 0.106 |
| *DENND1C* | chr19:6475456-6477198 | 7 | 3.2E-68 | 0.106 |
| *FMNL1* | chr17:43318045-43319382 | 7 | 2.0E-75 | 0.106 |
| *HNF4A* | chr20:43028501-43029997 | 9 | 4.6E-93 | 0.107 |
| *GATA6* | chr18:19756582-19758221 | 7 | 9.0E-64 | 0.107 |
| *HOXA6* | chr7:27180888-27185512 | 44 | 0.0E+00 | 0.108 |
| *HOXA3* | chr7:27152583-27156062 | 28 | 1.2E-234 | 0.108 |
| *LAMB3* | chr1:209825672-209825856 | 6 | 1.2E-76 | 0.108 |
| *MIR3193* | chr20:30195969-30196714 | 5 | 7.3E-53 | 0.108 |
| *APOH* | chr17:64225346-64226953 | 5 | 2.3E-34 | 0.108 |
| *DUSP6* | chr12:89747628-89749822 | 15 | 1.1E-108 | 0.109 |
| *PLGLA* | chr2:106959205-106959878 | 7 | 2.2E-75 | 0.109 |
| *RABGAP1L* | chr1:174843754-174844490 | 5 | 3.6E-51 | 0.109 |
| *HOXA-AS3* | chr7:27178861-27179432 | 5 | 2.1E-80 | 0.109 |
| *SFT2D3* | chr2:128453108-128453484 | 5 | 9.0E-19 | 0.109 |
| *FAAP20* | chr1:2120985-2121724 | 6 | 5.0E-34 | 0.109 |
| *PLEKHM3* | chr2:208794914-208795859 | 5 | 2.1E-49 | 0.109 |
| *IGF2BP1* | chr17:47090616-47092178 | 8 | 4.5E-59 | 0.110 |
| *SH2D3C* | chr9:130515119-130517848 | 6 | 2.7E-43 | 0.110 |
| *PDZK1* | chr1:145726979-145727762 | 6 | 6.9E-54 | 0.112 |
| *HOXA3* | chr7:27159883-27166103 | 18 | 2.0E-79 | 0.113 |
| *EIF2AK4* | chr15:40268421-40269214 | 5 | 8.0E-24 | 0.114 |
| *BMP4* | chr14:54418728-54420185 | 6 | 9.8E-21 | 0.115 |
| *ADGRG1* | chr16:57653169-57654347 | 5 | 2.8E-58 | 0.115 |
| *TRIO* | chr5:14405632-14406585 | 5 | 4.5E-13 | 0.116 |
| *TOLLIP* | chr11:1325718-1327450 | 10 | 3.2E-52 | 0.117 |
| *PSMG3* | chr7:1606266-1607787 | 8 | 6.3E-71 | 0.118 |
| *TNNC1* | chr3:52487733-52488229 | 5 | 6.1E-74 | 0.120 |
| *HLX* | chr1:221057236-221059414 | 8 | 2.5E-22 | 0.129 |
| *ZNF436* | chr1:23696021-23698143 | 8 | 5.8E-60 | 0.166 |
| *HLA-DPB1* | chr6:33046344-33049505 | 22 | 5.9E-204 | 0.167 |

**Figure 2.**
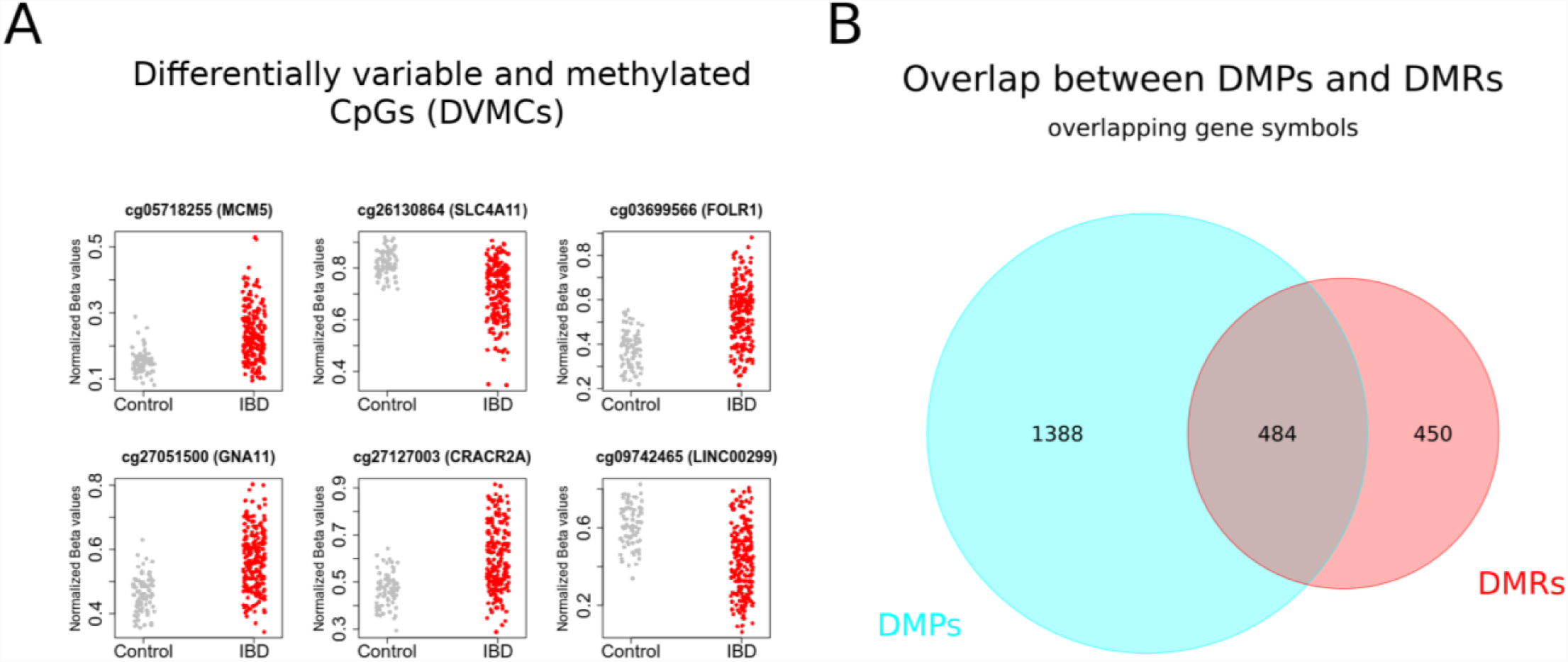
Mean DNA methylation and variability distinguishes IBD from healthy intestinal epithelial cells. **A.** Top differentially variable methylated CpG sites (DVMCs) in IBD *vs* Control. DNA methylation was plotted as beta values for each of the top nine DVMC identified with the iEVORA algorithm (see Methods section). **B.** Gene symbols overlapping between site-(DMPs) and region (DMRs)-level analyses (Representation factor: 5.5, p < 1e-5).

In addition to mean methylation differences at the probe and region levels (i.e. DMPs and DMRs), methylation variation has been associated with disease and cancer susceptibility ^23^. To explore this, we used the iEVORA algorithm in the same datasets, to identify differentially variable and methylated CpGs (DVMCs). Using stringent criteria of differential methylation and variation, we identified 4583 DVMCs (Fig 2A and Table S3). Of note, for most of these sites (71%), IBD samples displayed higher variability than control tissues.

In summary, the intestinal epithelia of IBD displays large non-random methylome abnormalities characterized by high variability, but also by absolute changes in mean DNA methylation at particular loci.

### Genomic and biological context of IBD-associated DNA methylation changes in intestinal epithelia

DMPs distinguishing IBD from control tissues were assessed for genomic distribution, in terms of gene-centric and CpG island (CGI)-centric context. DMPs were relatively absent from CGIs, gene promoters, or the vicinity of transcription start sites (TSS) (Fig 3A-3C). Instead, hypo and hypermethylated DMPs were highly concentrated in non-CGI regions (i.e. open sea) (Fig 3A).

**Figure 3.**
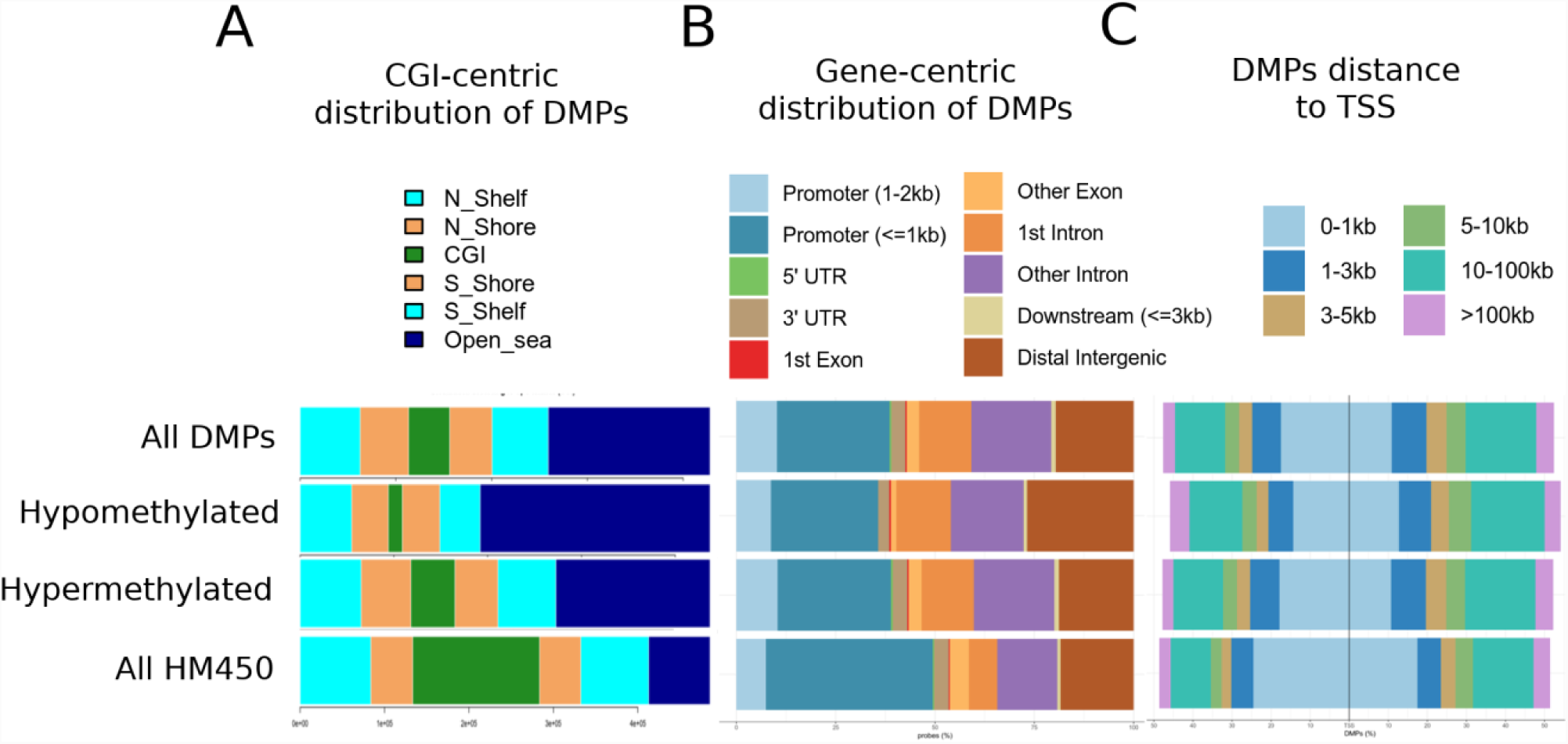
Genomic distribution of IBD-related DMPs. DMPs were annotated according to CpG islands (CGI) (**A)**, relation to gene features **(B)**, and distance to the nearest transcription start site (TSS) **(C).** For each genomic context, distribution is shown separately for all DMPs, those hypo or hypermethylated in IBD relative to healthy tissues, and all the HM450 probes, as a control.

Pathway analysis of DMRs revealed over-representation of pathways related to metabolism and signal transduction, including Adipogenesis, Hemostasis, G alpha signaling events, Pathways in cancer, and TGF-beta Receptor Signaling (Table 4).

**Table 4.** Pathway analysis.

| Pathway | Adjusted P-value | Combined Score | Dataset |
| --- | --- | --- | --- |
| *Adipogenesis genes_Mus musculus_WP447* | 5.78E-06 | 33.17 | WikiPathways 2016 |
| *Adipogenesis genes_Homo sapiens_WP236* | 5.78E-06 | 32.1 | WikiPathways 2016 |
| *TGF Beta Signaling Pathway_Mus musculus_WP113* | 4.28E-04 | 26.34 | WikiPathways 2016 |
| *TGF-beta Receptor Signaling_Homo sapiens_WP560* | 7.32E-04 | 24.45 | WikiPathways 2016 |
| *Alpha6-Beta4 Integrin Signaling Pathway_Mus musculus WP488* | 2.44E-03 | 18.52 | WikiPathways 2016 |
| *Hemostasis_Homo sapiens_R-HSA-109582* | 1.31E-03 | 29.15 | Reactome |
| *G alpha(12/13) signalling events_Homo sapiens_R-HSA-416482* | 4.29E-03 | 21.79 | Reactome |
| *Pathways in cancer_Homo sapiens_hsa 05200* | 5.90E-04 | 26.06 | KEGG 2016 |
| *Aldosterone synthesis and secretion_Homo sapiens_hsa04925* | 5.90E-04 | 24.68 | KEGG 2016 |

Overall, abnormal DNA methylation in IBD is relatively absent from CGIs. At the biological level, DNA methylation changes are enriched in inflammation-related pathways. Such changes may occur downstream of cytokine signaling. Alternatively, they may represent early changes linked to genetic susceptibility.

### IBD DMPs are genomically closer to IBD risk polymorphisms, and are enriched on blood mQTLs

DNA methylation may represent an intermediary between genotype and disease susceptibility, and such genetic influences on DNA methylation within a defined genomic context are known as methylation quantitative trait loci (mQTLs). Among DMRs with a significant genetic association, we found *ITGB2, MUC16, JAK3, KRT8*, and *HLA* genes, confirming the findings of previous studies ^7,24–26^. Moreover, some DMPs display a bimodal DNA methylation distribution (see Methods), suggesting that their methylation levels are directly dependent on genotype. To explore a genotype-methylation association, we calculated the genomic distance between DMPs identified in our analysis and single nucleotide polymorphisms (SNPs) associated with IBD risk ^24,25,27^. Of note, DMPs were overall significantly closer to a known IBD risk SNP, compared all HM450 sites taken together (Fig 4). This difference was preserved after independently comparing hyper or hypomethylated DMPs, and consistent across three independent SNP datasets (Fig 4 and S2C).

**Figure 4.**
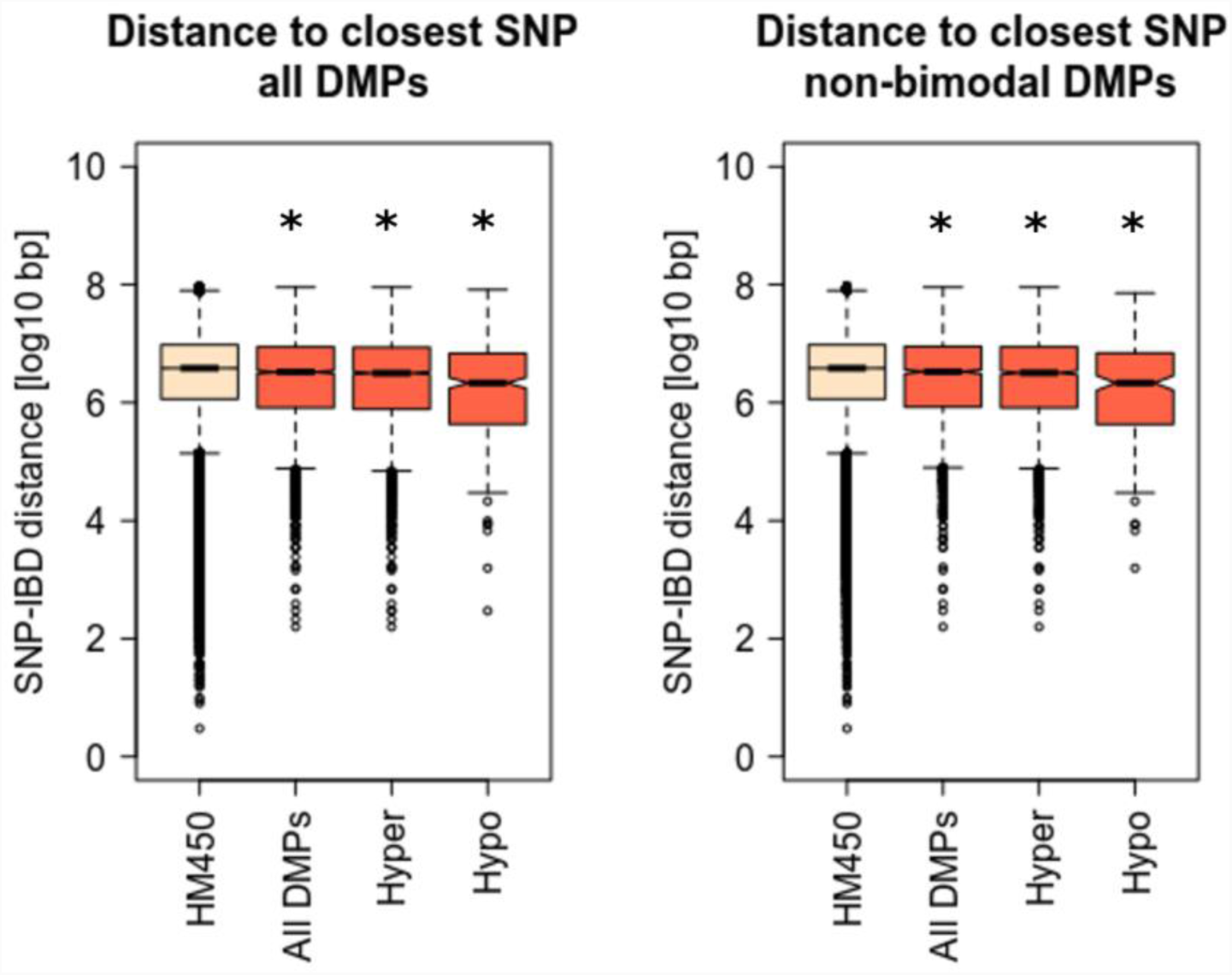
Genomic distances between IBD-related DMPs and known risk SNPs. Shortest genomic distances were calculated between each IBD-related DMP and the closest IBD-associated polymorphism (SNP). Boxplots represent the distribution of such distances for all DMPs, or separately for hyper- or hypo-methylated DMPs. The distance of all HM450 CpG sites was calculated as a control (left boxplot in both panels). The same analysis was performed for all DMPs (left panel) or using only DMPs that did not display a bimodal distribution (right panel), as described in Methods. (*) denotes a significant difference in mean distance relative to control HM450 distances (p < 1e-5).

Moreover, IBD-DMPs showed a slight but significant enrichment in CpGs participating to blood mQTLs (representation factor=1.1, p < 0.002) as defined by McRae et al. ^28^. In fact, 566 out of the 4280 DMPs participated to the 52916 mQTLs reported previously (Supplementary Table S4). To ascertain whether the SNPs putatively associated to our DMPs were also associated to IBD, we interrogated the largest fine-mapping study performed to date on the disease that claims to identify associations at a base-pair resolution level ^24^. We found that 4 of the 566 mQTLs identified here bear an IBD-associated polymorphism, namely rs11264305, rs17228058, rs3806308 and rs3807306, located in or close to *ADAM15, SMAD3, RNF186* and *IRF5*, respectively (Figure S3). Briefly, we found that SNP-CpG pairs overlap regulatory loci, discernible by H3K27ac histone marks and the presence of a CpG island (in the case of *ADAM15*).

These findings suggest that at least a fraction of IBD abnormal methylome is in direct relationship with upstream genetic susceptibility variants.

### IBD and epithelial and immune cell fractions of the celiac duodenum share DMPs

As the IBD methylome is both, related to inflammation and genetic susceptibility, it may also be largely unspecific. We therefore chose celiac disease (CeD), a chronic inflammatory condition of the GI tract with a well characterized genetic component, to get further insight into methylome specificity. In addition, DNA methylation data for epithelial and immune components of CeD were analyzed separately ^29^. When we crossed IBD-DMPs with epithelial CeD-DMPs we found that, out of 4280 IBD-DMPs and 43 CeD epithelial-DMPs, 7 were common (representation factor=17.1, p < 1e-05) (Table 5). Interestingly, 5/7 common DMPs mapped to the HLA region on chromosome 6. On the other hand, 31 IBD-DMPs were common with the 310 CeD immune-DMPs (representation factor=10.5, p < 1e-5). These common hits were enriched for TGF-β signaling pathway (WikiPathways, adjusted p value=0.04419), and were spread across the genome. All common DMPs followed the same direction (i.e. hypo or hypermethylation) in both diseases, indicating that methylation alterations were concordant. However, methylation fold changes were larger in CeD, probably due to the fact that the celiac DMPs were identified in separated cell populations, while IBD methylation was assessed in whole intestinal tissue potentially blurring cell-specific signatures.

**Table 5.**
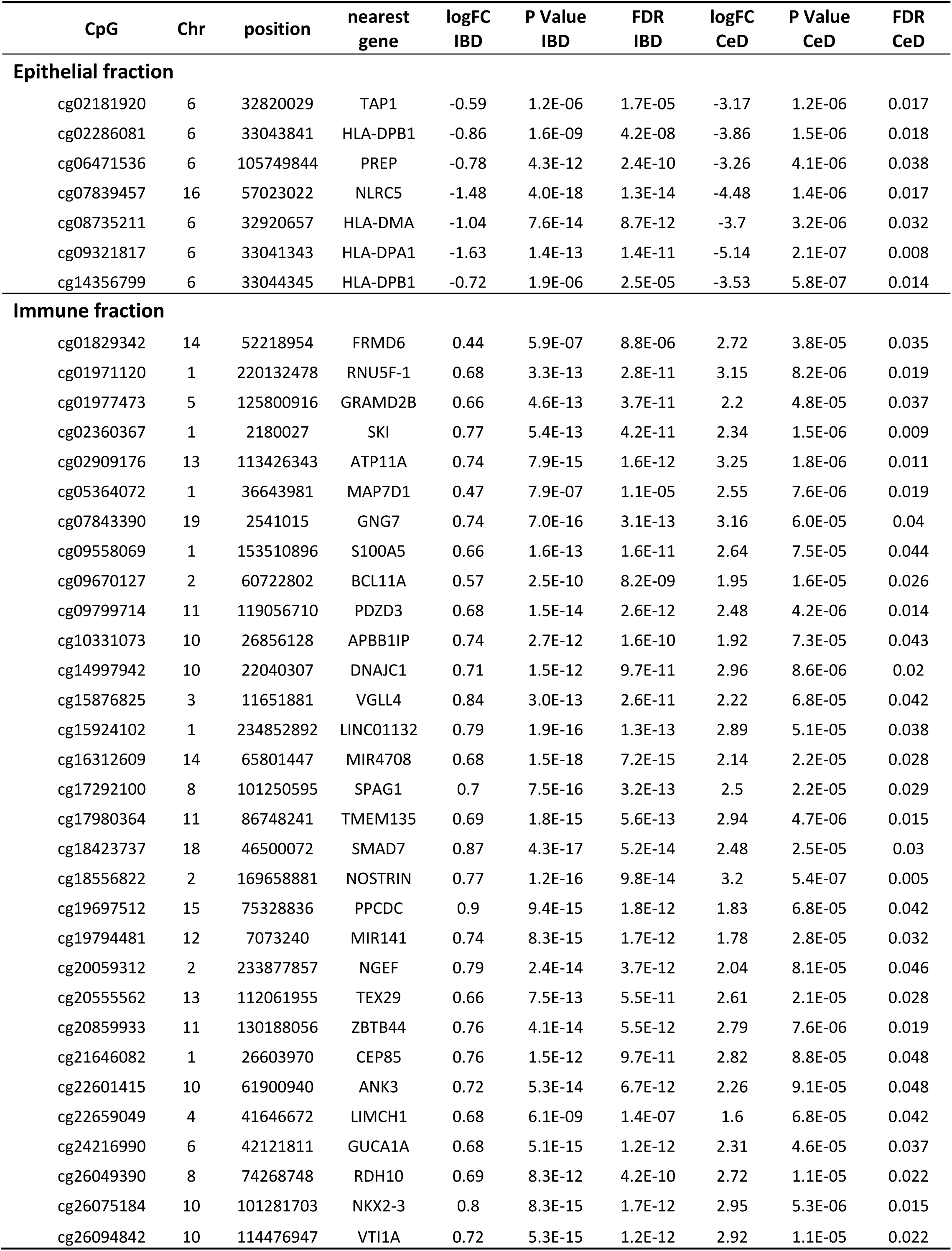
IBD DMPs previously identified to be differentially methylated in both CeD duodenal epithelia and immune fractions.

In summary, there is a significant overlap in DNA methylation changes associated with IBD and CeD, including the HLA region.

## Discussion

IBD is a complex pathology with a wide range of clinical trajectories. Despite such heterogeneity, we show here that methylome-wide changes in IBD are robust to main clinical parameters and consistent across several studies. Non-random changes in mean methylation and increased methylation variability are characteristic of IBD. Although an increased variance in DNA methylation values could be an unspecific effect of inflammation, higher methylation variability has also been described in the vicinity of tumoral tissues ^23^. This may be of interest, considering that a fraction of IBD subjects have an increased risk of colon cancer ^6^.

Different characteristics of DNA methylation, such as its relative stability, make this mark an ideal sensor of disease risk and progression. However, a deeper mechanistic insight is necessary to better distinguish those methyl marks that are dependent on genetic susceptibility from those that are a consequence of environmental cues. We suggest here that IBD methylome is indeed a combination of both components: on the one hand, many associations at the site and region levels were enriched in inflammatory pathways, suggesting that methyl marks could have been introduced downstream of cytokine signaling. On the other hand, at least a fraction of DNA methylation changes were linked to a neighboring risk polymorphism, indicating an effector role for DNA methylation in the interface between genotype and phenotype.

In agreement with the largest study selected for our meta-analysis ^9^, genes near abnormal DNA methylation were enriched in immune and inflammatory pathways, highlighting the role of chronic inflammation in both, UC and CD. In particular, TGF-β is a cytokine able to modulate the inflammatory response, and it was enriched in IBD-DMRs. Moreover, it was enriched in those DMPs common between IBD and CeD, in agreement with the crucial role of TGF-β pathway in regulating the intestinal T cell response. An additional element that emerged from our pathway analysis is the potential crosstalk between IBD and adipogenesis. In fact, patients with IBD, particularly those with CD, develop ectopic adipose tissue (fat-wrapping or creeping-fat) covering a large part of the small and large intestine ^30^. It has been proposed that in obese or overweight IBD patients it is the mesenteric adipose tissue that contributes to intestinal and systemic inflammation ^30^.

In terms of genomic distribution, we found that DMPs are relatively absent from CGIs. Instead, they could be associated with other regulatory regions such as enhancers, for example in association with SNPs. Indeed, GWAS performed in multiple complex diseases have shown that SNPs of susceptibility are enriched in enhancer regions, and DNA methylation could be an intermediary in this process ^31,32^. Illustrating this, the presence of differentially methylated sites in the vicinity of known susceptibility loci supports the notion of DNA methylation as an intermediary between genotype and phenotype (mQTLs). In addition, among DMRs with a significant genetic association, we find *ITGB2, MUC16, JAK3, KRT8, HLA* genes, all of them associated with a role in IBD pathogenesis ^33–37^.

The enrichment of CpGs participating in both IBD-DMPs as well as mQTLs suggests that a considerable number of the DMPs identified in our metanalysis are regulated by SNP-genotypes in cis. However, very few of these are associated with IBD. This observation points to the possibility that, although fine-mapping aims to identify the SNPs responsible of the disease-association, other nearby SNPs in strong linkage disequilibrium could be the ones implicated in the mQTLs, drawing the methylation patterns reported. Additionally, we describe a picture in which most of the IBD-DMPs seem to be genotype-independent, since they do not participate in any mQTL, at least in blood. Regarding the SNPs associated to IBD as well as to the methylation levels of IBD-DMPs, it is interesting that the methylation of a CpG island 4 kb upstream of the cg24032190-DMP identified in the first intron of *SMAD3* has been reported to be allele-specific and to regulate the expression of the gene ^38^. Therefore, we propose another DMP in the same region that could mediate the association between the locus and IBD; and hypothesize that this could also be the case for the genomic regions surrounding *ADAM15, RNF186* and *IRF5*.

Regarding celiac epithelial DMPs also found altered in IBD, it is important to note that most of them were located in the HLA region. This locus presents strong linkage disequilibrium and encodes a number of genes related to immune response and immune regulation through self-recognition ^37,39^, and strongly predisposes to autoimmune diseases such as CeD. In our previous work ^29^, we claimed to have found a genotype-independent methylation signature in celiac duodenal epithelia. The finding of a signature in the HLA region common to IBD and CeD reinforces this idea, given that the HLA association with IBD is much weaker (variance explained <5%) than with CeD, and moreover, different HLA haplotypes drive these associations ^33^. Additionally, this common methylation signature points to a non-specific pattern, probably responding to common inflammatory forces in the two disorders.

## Conclusions

Our findings illustrate an aberrant DNA methylation landscape in IBD, independent of IBD subtype and other clinical and pathological features. The enrichment of abnormal DNA methylation in inflammatory pathways and genes suggests a direct role for this mark downstream of cytokine signaling and/or a risk genotype. Such a landscape may be a more general indicator of intestinal chronic inflammation.

## Methods

### Dataset Selection

Dataset selection criteria included: methylome data obtained from intestinal mucosa (including colon and terminal ileum), availability of healthy controls and IBD samples (CD, UC, or both), in data obtained using Human Infinium Bead Arrays (Illumina’s HM450 or EPIC arrays), an established technology to detect DNA methylation ^40^. Table 1 shows the main characteristics of the datasets fulfilling these criteria.

### Data Preprocessing

All methylation data and sample information were downloaded from Gene Expression Omnibus (GEO) and Array Express public repositories, and analyzed using R/Bioconductor packages ^41^. Normalized data was loaded into R directly from each repository, except when raw idat files were also available. In that case, idat files were normalized using the “Funnorm” function of the minfi package ^42^. Each dataset was independently assessed for data quality and distribution, before merging. Merged data was filtered for sex chromosomes, known cross-reactive probes ^43^, and probes associated with common SNPs that may reflect underlying polymorphisms rather than methylation profiles ^44^. In addition, the “nmode.mc” function of the ENmIx package was used for the identification of multimodal sites ^45^. These sites were not removed at this step, but were used instead to classify significant associations in a later step.

### Quality Control and Batch Correction

After filtering, 393112 CpG sites common to all datasets were used to identify principal components (PC) of variation and plotted using PC regression and multidimensional scaling (MDS) plots. Strong associations were observed between PCs and known variables (i.e. dataset, sex, age, and anatomical location), with age and anatomical location partially confounded by the dataset of origin. Latent variables were also identified, using surrogate variable analysis ^46^. As additional quality control, DNA methylation values were used to predict age and sex and contrast with downloaded phenotype information. Sex was inferred from the median total intensity signal on XY chromosomes, and permitted the identification of 8 sex mismatches that were removed from the analysis (Fig S1). Age prediction was performed using Horvath’s coefficients ^47^, as implemented in the wateRmelon package ^48^. There was an overall correlation between reported and predicted age (Fig S1). For two datasets where age was not available, predicted age corresponded to adult samples, as reported in the corresponding repositories. The common merged and filtered matrix of methylation beta values and their corresponding phenotype data was taken to the next step.

### Differential Methylation

Associations were tested for 393112 CpG sites, across 285 samples (81 control and 204 IBD samples). Methylation data was modeled at the probe and region levels using a linear model with Bayesian adjustment ^49^. Sex and dataset were modeled together with subject status (i.e. control or IBD patient). Surrogate variables identified in the previous step were also included in the linear model to account for unknown sources of variation. Quantile-quantile (QQ) plots were used to inspect the distribution of resulting p values and estimate statistical inflation. Differentially methylated positions (DMPs) and regions (DMRs) were selected based on a methylation change (delta beta) of at least 10% or 5% (for DMPs and DMRs, respectively) when comparing control vs. IBD samples, and a false discovery rate-(FDR) adjusted p value below 0.05. DMRs were identified with the DMRcate package using the recommended proximity-based criteria ^50^. A DMR was defined by the presence of at least two differentially methylated CpG sites with a maximum gap of 1000 bp. To identify CpG positions exhibiting significant differential variation and differential methylation (DVMCs), data was analysed using iEVORA, an algorithm that identifies DNA methylation outlier events shown to be indicative of malignancy ^51^. iEVORA is based on Bartlett’s test (BT) that examines the differential variance in DNA methylation, but because BT is very sensitive to single outliers, it is complemented with re-ranking of significant events according to t-statistic (TT, t test), to balance the procedure. The significance is thus assessed at the level of differential variability, but the significance of differential variability with larger changes in the average DNA methylation are favored over those with smaller shifts. We used adjusted q(BT) <0.001 and p(TT) <0.05 as thresholds for significant DVMCs. To study genomic context, we used HM450 annotations, with hg19 as the human reference genome, UCSC and previously reported genomic features ^52^. Differentially methylated genes (DMPs, DMRs, and DVMCs) were further analyzed to determine functional pathways and ontology enrichment using Enrichr ^44^.

### SNPs-DMPs associations in IBD and CeD

To identify methylation quantitative trait loci (mQTL), single nucleotide polymorphisms (SNPs) associated with IBD risk were obtained from a fine-mapping study of IBD with single-variant resolution ^24^. Two independent GWAS were also considered in some of the analyses: 1. Jostins L et al.^27^, and 2. Lange KM de et al.^25^. The genomic distances between 368 unique SNPs pooled from these three studies and IBD-associated DMPs were calculated using the R package GenomicRanges. In addition, we searched for those CpGs that apart from being differentially methylated in IBD according to our metanalysis, were previously reported to be differentially methylated in a previous work performed by our group in CeD ^29^. CeD is a genetic, inflammatory condition of the duodenum in which the Human Leucocyte Antigen (HLA) region explains around 40% of the heritability, and HLA-DQ2/-DQ8 molecules are necessary for gliadin presentation and activation of the autoimmune response. Briefly, we looked for the overlap between the bimodal IBD-DMP list presented here and the celiac DMPs found in both the epithelial and the immune cell fractions of the duodenum. We also searched for the IBD-DMPs that were previously reported to participate in blood mQTLs in cis (2 Mb, p < 1e-6), according to the largest to-date mQTL database available ^28^, and found the overlap between them and the SNPs associated to IBD ^24^. All the overlaps were reported using in-house R scripts. We also calculated the representation factor and the associated probability of the overlaps (http://nemates.org/MA/progs/overlap_stats.html), in order to stablish whether they were significant.

**Abbreviations**
CD: Crohn’s disease
CeD: celiac disease
CGI: CpG island
DMP: differentially methylated position
DMR: differentially methylated region
DVMC: differentially variable and methylated cytosine
GI: gastrointestinal
GWAS: genome-wide association studies
HLA: human leukocyte antigen
IBD: inflammatory bowel disease
mQTL: methylation quantitative trait loci
SNP: single nucleotide polymorphism
TGF-β: transforming growth factor beta
UC: ulcerative colitis

## Authors’ contributions

IA and HH carried out the methylation data meta-analysis; NF and JRB performed the mQTL analyses and IBD-CeD comparison; IA, HH, RD and CG co-wrote the first draft of the manuscript; JM and HH conceived the study. All authors read, critically revised and approved the final manuscript.

## Acknowledgements

We thank the patients involved in the research and all researchers that deposited their data in open repositories. This work was supported by the Agence Nationale de Recherches sur le SIDA et les Hépatites Virales (ANRS, Reference No. ECTZ47287 and ECTZ50137), the Institut National du Cancer AAP PLBIO 2017 (project : *T cell tolerance to microbiota and colorectal cancers*), and La Ligue Nationale Contre Le Cancer Comité d’Auvergne-Rhône-Alpes AAP 2018. NF is partially funded by the Basque Department of Health (project 2018/111086 to JRB).

## Supplementary Data

**Table S1.** Full list of differentially methylated positions (DMPs).

**Table S2.** Full list of differentially methylated regions (DMRs).

**Table S3.** Full list of differentially variable and methylated CpGs (DVMCs).

**Table S4.** Overlap between bimodal IBD-DMPs and CpGs participating to blood mQTLs as defined by McRae et al. ^28^.

**Figure S1.**
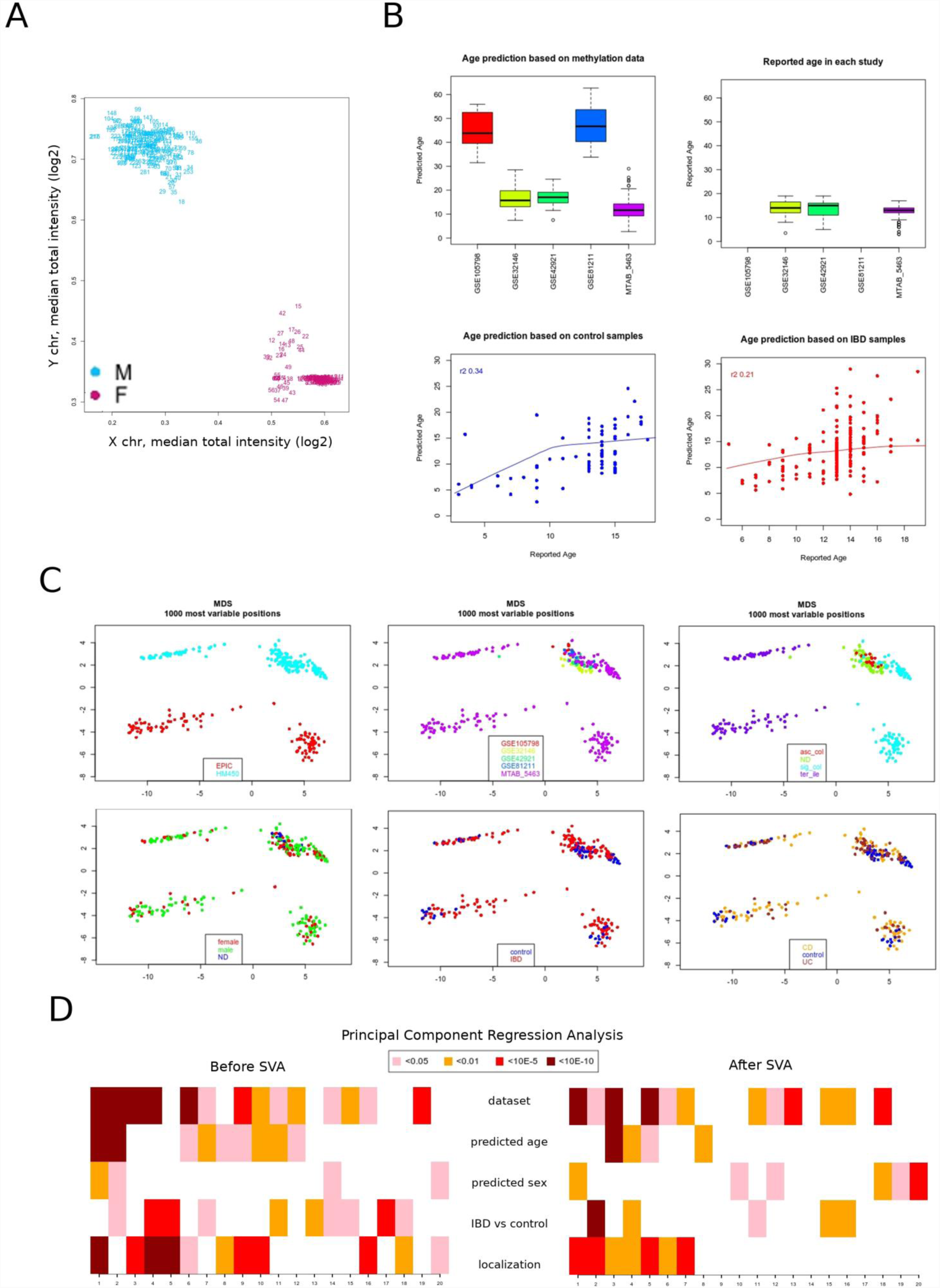
Data quality and preprocessing. **A.** Sex predictions using XY chromosome methylation data. **B.** Predicted (upper left panel) and reported (upper right panel) age. Reported age was not available for the two datasets that studied adult subjects. Matching between reported and predicted age in control and IBD samples (lower left and right panels, respectively). **C.** Multidimensional scaling (MDS) plots, according to bead array version (EPIC vs HM450), dataset, sex, condition (control vs IBD), anatomical location (asc_col: ascending colon, sig_col: sigmoid colon, ter_ile: terminal ileum, ND: no data available), and IBD subtype (CD vs UC). **D.** Principal component regression analysis (see Methods) before (top) and after (bottom) adjustment.

**Figure S2.**
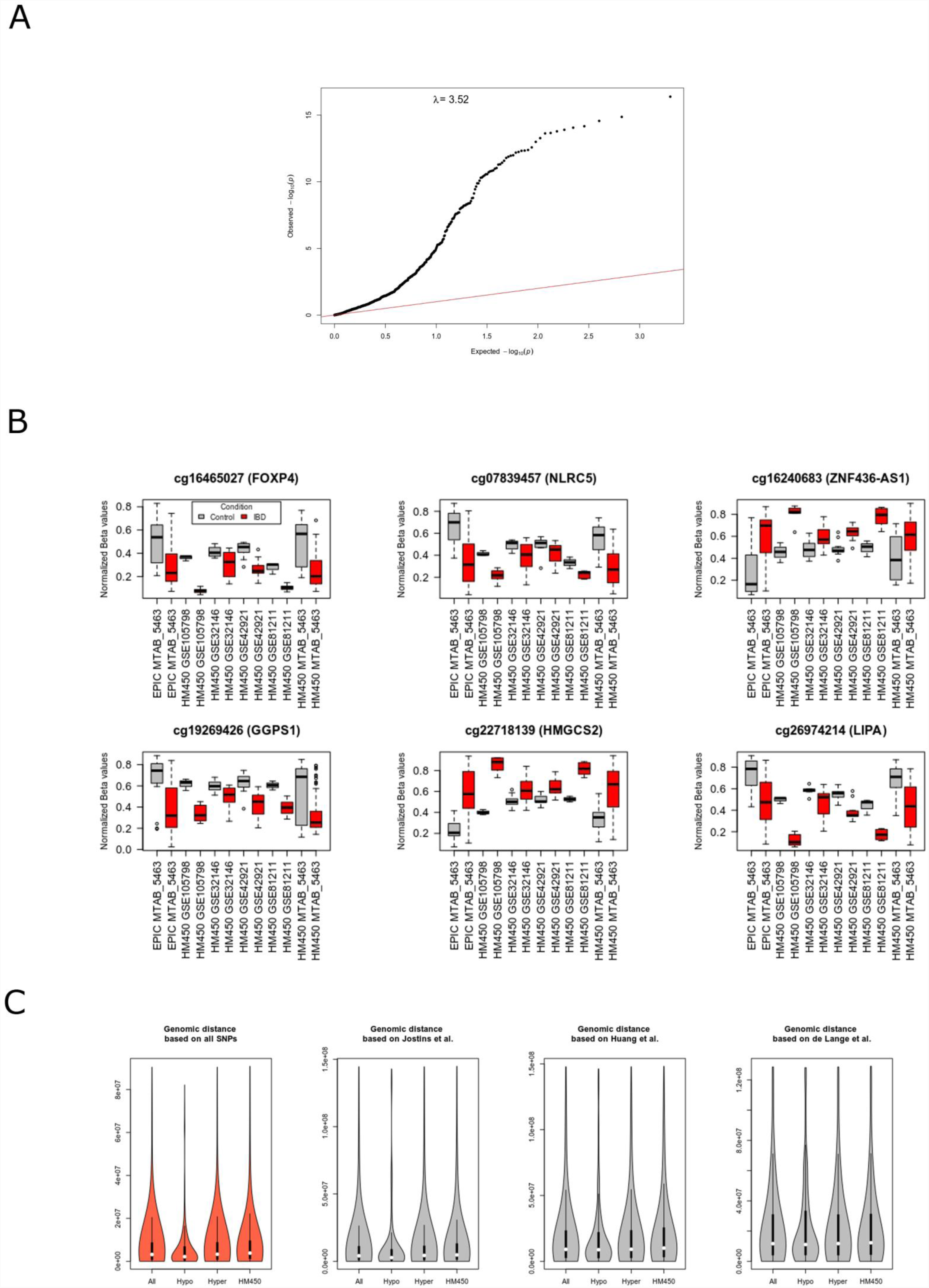
Differential methylation analysis was perfomed using a linear regression model (see Methods). **A.** Quantile-quantile plot for the association between DNA methylation at the probe-level and sample type (IBD vs control). Statistical inflation (λ) is indicated on the plot. **B.** Distribution of the top DMPs across the different datasets (compare with Fig 1A). Barplots are shown for control (gray) and IBD (red) samples separately for each dataset. **C.** Genomic distances were calculated between all DMPs identified and known SNPs associated with IBD risk, according to three independent studies ^24,25,27^. Left violin plot shows the distribution of the distances using the aggregated data from the three studies (red). The remaining three plots show the distances independently for each of the studies. All: all DMPs, Hypo: DMPs hypomethylated in IBD, Hyper: DMPs hypermethylated in IBD, HM450: all informative CpG sites in the Infinium bead array.

**Figure S3.**
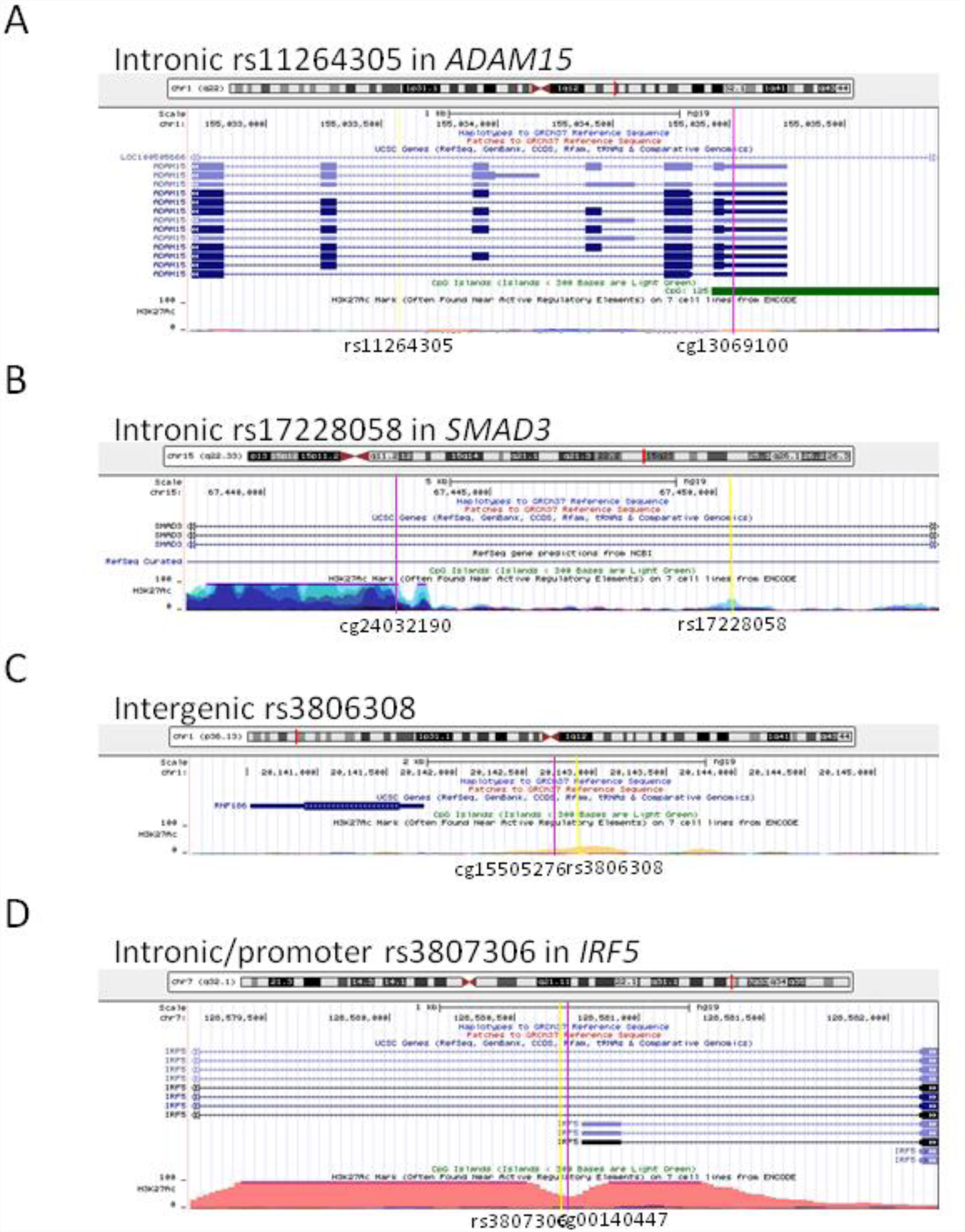
Genomic locations of the IBD-associated SNPs rs11264305, rs17228058, rs3806308 and rs3807306. These SNPs participate to mQTLs according to McRae et al. together with the IBD-DMPs cg13069100, cg24032190, cg15505276 and cg00140447, respectively. SNPs are marked by yellow lines, while location of CpG sites is highlighted by pink lines. CpG islands as well as layered H3K27ac histone marks are also depicted as indicated in the Y axis (based on the UCSC Genome Browser, https://genome.ucsc.edu/).

